# JNK MAPK Regulates IFN-Stimulated Genes and Cell Adhesion in Chemoresistant, Quiescent Leukemic Cells

**DOI:** 10.1101/689570

**Authors:** Sooncheol Lee, Alireza Ameen Baradar, Paola A. Sundaram Buitrago, Elijah Puente, Zeynep Kilinc, Olivia Cox, Shobha Vasudevan

## Abstract

Cancer cells survive environmental stresses including DNA damage-inducing chemotherapy by entering cellular quiescence (G0). In our previous study, to elucidate how G0 leukemic cells become resistant to chemotherapy, we profiled them at the transcriptome, translatome and proteome levels. We interrogated these datasets and find that IFN-stimulated genes are enhanced in our G0 models of leukemic cells induced by chemotherapy or serum-starvation. Mechanistically, upon induction of G0, STAT1 is phosphorylated on tyrosine 701, leading to transcription of IFN-stimulated genes. Importantly, our data reveal that activation of JNK and p38 MAPK is integral to STAT1 phosphorylation and expression of IFN-stimulated genes. Pharmacological inhibition of JNK or p38 MAPK greatly reduced expression of IFN-stimulated genes as well as STAT1 phosphorylation, revealing a JNK-STAT1 pathway that regulates IFN-stimulated genes. We find that the JNK-STAT1 pathway enhances adherence of G0 leukemic cells. Consistently, inhibition of the JNK-STAT1 pathway dramatically reduced the number of adherent G0 cells. STAT1 is transiently phosphorylated within 24 hours of chemotherapy, leading to transcriptional expression of IFN-stimulated genes. Subsequently, translational regulation attenuates their expression in G0 leukemic cells. These studies uncover translational and transcriptional regulation of IFN-stimulated genes by a JNK-STAT1 pathway that enhances cell adhesion in G0 leukemic cells.

## Introduction

Cellular quiescence (G0) can be induced under unfavorable conditions such as chemotherapy, serum-starvation, high confluency and contact inhibition [1–3]. In our previous study, G0 acute monocytic leukemic (AML) cells were induced by treatment of proliferating THP1 cells and other AML cell lines with AraC (cytosine arabinoside) chemotherapy or serum-starvation [4]. These G0 cells are resistant to subsequent AraC treatment and re-enter cell cycle under favorable conditions [4] (Fig. 1A). To understand how G0 leukemic cells survive chemotherapy, their transcriptome, translatome (polysome-associated RNAs) and proteome were systematically analyzed over time of treatment [4]. These studies revealed that gene expression in our G0 models of leukemic cells recapitulates that of *in vivo* chemoresistant leukemic models [4]. In this study, we used these datasets to identify differentially expressed genes in chemoresistant G0 leukemic cells.

**Figure 1.**
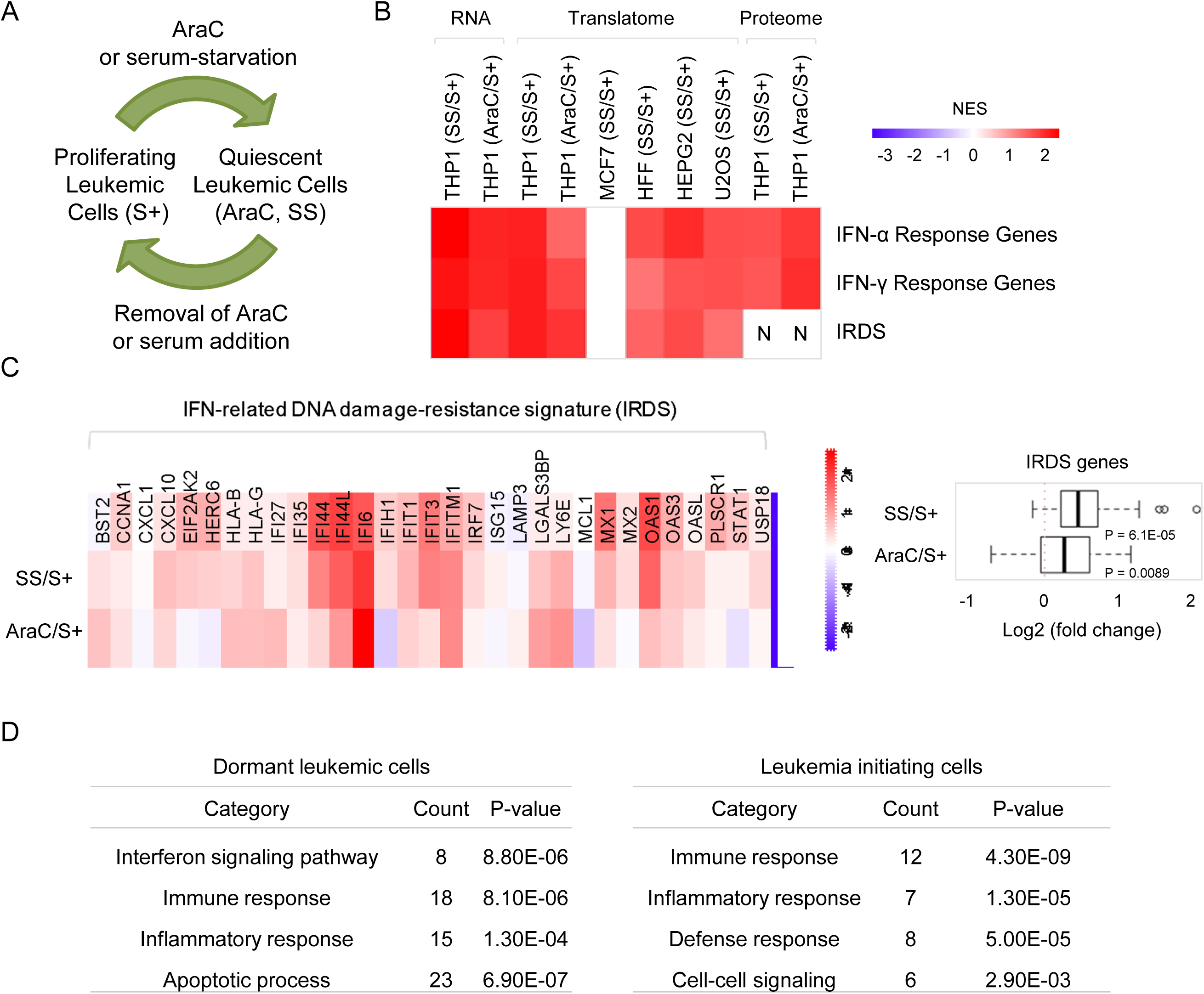
Expression of IFN-stimulated genes, including IRDS genes, is augmented in G0 leukemic cells. **A.** Induction of G0 leukemic cells (referred to as AraC and SS cells based on treatment). Treatment of proliferating leukemic cells (referred to as S+ cells) with 5 μM AraC or serum-starvation for 3-4 days, arrests the cells in the G0 phase and renders them resistant to subsequent AraC treatment [4]. When AraC is removed or serum is added, AraCS and SS cells restart proliferation. **B.** GSEA analysis of IFN-stimulated genes including IRDS genes, in the transcriptome, translatome and proteome datasets of G0 and proliferating cancer cells. Normalized enrichment scores are shown in color code. Inapplicable analyses are marked with ’N’. **C**. Expression of IRDS genes in the translatome level in G0 leukemic cells is shown as a heatmap (left) or a boxplot (right). **D**. DAVID gene ontology analyses of signature genes of dormant leukemic cells and leukemia initiating cells.

Surprisingly, we find that interferons (IFNs)-stimulated genes were significantly increased in G0 leukemic cells compared to proliferating or untreated leukemic cells. IFN cytokines are induced by multiple stress signals such as viruses and other stressors, and bind to cell surface receptors and initiate a signaling cascade through the Janus kinase/Signal transducer and activator of transcription (JAK-STAT) pathway, producing IFN-stimulated genes [5–7]. Although IFN-stimulated genes are traditionally believed to act as antiviral mediators [5, 6, 8, 9], many studies show that a subset of IFN-stimulated genes can impact survival of cancer cells under genotoxic stress via tumor-intrinsic and -extrinsic mechanisms [10–12]. IFN-stimulated genes are induced by transcriptional pathways triggered by IFNs, are diverse, and impact both innate immunity and adaptive immunity and can control the anti-tumor immune response [12]. IFNs appear to have dual impact on tumors, where they can promote tumor persistence and yet also can improve response to therapy including immunotherapy [13, 14]. Cancer cells promote type I IFN (IFNα, IFNβ) synthesis but inhibit their own ability to respond to IFN, thus enabling the pro-tumor IFN-related DNA damage resistance signature (IRDS) response while the response to the cytotoxic IFN type I effects is inhibited [15]. Type II (IFNγ) -stimulated gene signature is associated with efficiency of immunotherapy response; however, IFNγ is also associated with enhanced metastasis, and immune escape by tumors [11]. Several studies also demonstrate such dual-edged roles of IFN in acute myeloid leukemias, where IFN production enables therapeutic efficacy [16–18] but also enables immune suppression and poor prognosis [19]. For e.g.; IFNγ release by in vivo AML modifies bone marrow stromal cell gene expression to create an immunosuppressive environment [20]. While IFN-stimulated genes have powerful roles in many cancers, our findings show that G0 chemoresistant THP1 AML cells enhance both types of IFN-stimulated genes compared to untreated THP1 cells. How IFN-stimulated genes are regulated and what their biological functions in chemoresistant leukemic cells are, remain to be fully uncovered.

In this study, we find that the expression of IFN-stimulated genes is transcriptionally and transiently up-regulated *via* STAT1 phosphorylation. Subsequently, they are translationally down-regulated in AraC resistant leukemic cells. We find that the activation of JNK and p38 MAPK is essential for STAT1 phosphorylation and the expression of IFN-stimulated genes. Interestingly, the JNK-STAT1 pathway promotes cell adhesion of G0 leukemic cells; consequently, inhibition of the JNK-STAT1 pathway reduces cell adhesion of G0 leukemic cells. G0 leukemic cells also increase immune cell migration; however, abrogation of the STAT1 pathway did not diminish immune infiltration, indicating additional regulation in G0 leukemic cells to alter the tumor immune environment. We found a subset of the immunoglobulin superfamily and cell adhesion genes that are highly expressed in G0 leukemic cells, are suppressed upon STAT1 inhibition, indicating that these genes are targets of IFN-STAT1 pathway, and can contribute to cell adhesion in resistant G0 leukemic cells. Together, our data reveal insights into chemoresistant, G0 leukemic cells that induce immune cell migration, and uncover translational and transcriptional regulation of IFN-stimulated genes by a JNK-STAT1 pathway that enhances cell adhesion in G0 leukemic cells.

## Materials and Methods

### Cell Culture

THP1, MOLM13, U937 cells were maintained in RPMI 1460 media (Gibco) with 10% heat-inactivated FBS (Gibco), 2 mM L-Glutamine, 100 μg/ml streptomycin and 100 U/ml penicillin at 37°C in 5% CO_2_. MCF7 cells were grown as described previously [4]. To prepare resistant G0 leukemic cells, THP1 cells were washed with PBS followed by serum-starvation for 4 days or treated with 5 μM AraC for 3 days.

### Inhibitors and antibodies

AraC, Ruxolitinib [21], LY2228820 [22], BIRB796 [23], and JNK-IN-8 [24] were obtained from Selleckchem. Antibodies against Tubulin (05-829) and Actin (ABT1485) were from Millipore. Antibodies against IFIT2 (12604-1-AP), ISG15 (15981-1-AP), RPL13A (14633-1-AP), RPL11 (16277-1-AP), RPL17 (14121-1-AP) and Histone H3 (15303-1-AP) were from Proteintech. Antibodies against P-STAT1 (9167), STAT1 (9172), P-JNK (9251), JNK (9252), P-p38 MAPK (4511), p38 MAPK (9212) and IFIT1 (14769) were purchased from Cell Signaling Technology.

### Western blot analysis

Cells were lysed the buffer containing 40 mM Tris-HCl (pH 7.4), 6 mM MgCl_2_, 150 mM NaCl, 0.1% NP-40, 1 mM DTT and protease inhibitor cocktail (Roche). Cytoplasmic and nuclear fractions were separated as previously described [25]. About 80 μg of protein were loaded on SDS-PAGE gels (Bio-Rad) and transferred to PVDF membranes (Bio-Rad) as previously described for western analysis [4, 26].

### MTS and caspase 3/7 assays

Cell viability and caspase 3/7 activity were measured using Non-Radioactive Cell Proliferation Assay kit (Promega) and Caspase-Glo® 3/7 assay kit (Promega). A volume of 100 μL cells was placed in a 96-well plate, followed by addition of 20 μL MTS or 100 μL Caspase-Glo reagents. Absorbance at 490 nm and luminescence were measured after incubation.

### Cell adhesion assay

A 24-well plate was coated with 5 μg/ml human fibronectin (BD Biosciences) for 2 hours at 37°C. Proliferating and G0 THP1 cells were washed PBS and resuspended in media with 10% FBS. Cells were added into a 24-well plate at a density of 1 × 10^5^ cells / well and incubated for 2 hours at 37 °C in 5% CO_2_. The plate was washed with PBS to remove nonadherent cells and cells were stained with 0.2% crystal violet in 10% ethanol for 10 minutes. Microscopy images were taken and the number of adherent cells on the plate was determined.

### Cell migration assay

Transwell chambers (8 μm pore, Corning) were pre-equilibrated with serum-free media for 1 hour. GFP-tagged THP1 or MCF7 cells (2 x 10^4^/chamber) were placed in the top chamber, and 700 μL of proliferating or serum-starved THP1 cells containing media, in the bottom chamber. The bottom chamber with serum-starved THP1 cells was supplemented with 10% FBS as a control. The chambers were incubated at 37°C for 4 hours in 5% CO_2_. Cells on the upper surface of the filter were removed with a cotton swab. Migrated MCF7 cells on the underside of the filter were fixed in formaldehyde for 10 minutes and subsequently stained with 0.2% crystal violet. Migrated THP1 cells tagged with GFP were observed in the bottom chamber and visualized using a microscope. Microscope images were taken and the numbers of migrated cells were determined.

### Statistical analyses

Each experiment was repeated at least 3 times. No statistical method was used to pre-determine sample size. Sample sizes were estimated on the basis of availability and previous experiments [27, 28]. No samples were excluded from analyses. P values and statistical tests were conducted for each figure. Statistical analyses were conducted using Excel. SEM (standard error of mean) values are shown as error bars in all figures. Means were used as center values in box plots. P-values less than 0.05 were indicated with an asterisk. E-values were used for the statistical significance in the motif analysis.

## Results

### IFN-stimulated genes are significantly increased in resistant G0 leukemic cells

To determine expression level of IFN-stimulated genes in G0 cells, gene set enrichment analysis (GSEA) [29] was conducted on our previously acquired transcriptome, translatome and proteome datasets of G0 leukemic cells that were induced by treatment of proliferating THP1 cells with 5 μM AraC or serum-starvation over time for up to 3-4 days. Importantly, both IFN-α and IFN-γ stimulated genes are significantly up-regulated at the transcriptome, translatome, and proteome levels in G0 leukemic cells induced by AraC or serum-starvation, compared to proliferating cells (Fig. 1B, Table S1). Furthermore, different cell types induced to G0 by serum-starvation in HFF, HEPG2 and U2OS cell lines but not in the MCF7 cell line, increase expression of IFN-stimulated genes. Chemotherapy or radiotherapy induces expression of a subset of IFN-stimulated genes called the IFN-related DNA damage signature (IRDS) in breast cancer cells [30]. The expression level of this subset is significantly correlated with therapy resistance [30] in different types of solid tumors [30–42]. We found that IRDS genes are also highly expressed in G0 cells (Fig. 1B) including G0 leukemic cells (Fig. 1C). To test if high expression of IFN-stimulated genes in our G0 models of leukemic cells is conserved in *in vivo* resistant models of leukemia, we examined published transcriptome profiles of dormant leukemic cells (LRC) [43] and leukemia-initiating cells (LIC) [44] using the DAVID classification tool [45]. We found that interferon signaling pathway and immune response genes are significantly enriched in signature genes of LRC and LIC, respectively (Fig. 1D). These results confirm that our G0 leukemic cells are appropriate models to study how IFN-stimulated genes are regulated and their biological impact in resistant leukemia.

### IFN-stimulated genes are transcriptionally induced by STAT1 phosphorylation in resistant G0 leukemic cells

We asked how IFN-stimulated genes are expressed in resistant G0 leukemic cells. Mammalian cells release IFNs, a class of cytokines that trigger antiviral defense, modulate immunity, and impact tumor persistence via tumor-intrinsic and -extrinsic mechanisms [5, 6, 8–12]. IFN binding to its receptors phosphorylates and activates the STAT transcription factor family *via* JAK. Activated STATs dimerize and translocate into the nucleus and initiate transcription of IFN-stimulated genes [46]. To test if STAT proteins are activated, we examined them in a time course after G0 induction by Western blot analysis. We found that both AraC and serum-starvation induced STAT1 phosphorylation on tyrosine 701 in THP1 cells (Fig. 2A). These conditions specifically induce STAT1 activation as STAT3 or STAT5 were not phosphorylated (data are not shown). Interestingly, quantification of phosphorylated STAT1 in Fig. 2A shows that STAT1 was transiently phosphorylated at early times within 4-24 hours of serum-starvation or AraC treatment (Fig. 2B). In multiple leukemic cell lines, STAT1 was also phosphorylated on tyrosine 701 with 24 hours of AraC treatment (Fig. 2C). To investigate the localization of phosphorylated STAT1 in G0 leukemic cells, nuclear and cytoplasmic fractions were separated. As expected, phosphorylated STAT1 is distinctly localized to the nucleus one day after AraC treatment (Fig. 2D). To measure the effect of STAT1 phosphorylation on expression of IFN-stimulated genes, we analyzed the transcriptome of G0 leukemic cells upon inhibition of STAT1 phosphorylation. STAT1 phosphorylation was pharmacologically inhibited by the JAK inhibitor, ruxolitinib that is used to treat myelofibrosis [18, 21, 47–49] (Fig. 2E). We found that IFN-α, IFN-γ-stimulated genes including IRDS genes were significantly down-regulated upon ruxolitinib treatment, compared to vehicle (Fig. 2F) in AraC-treated cells. These data indicate that phosphorylation of STAT1 is responsible for the expression of IFN-stimulated genes.

**Figure 2.**
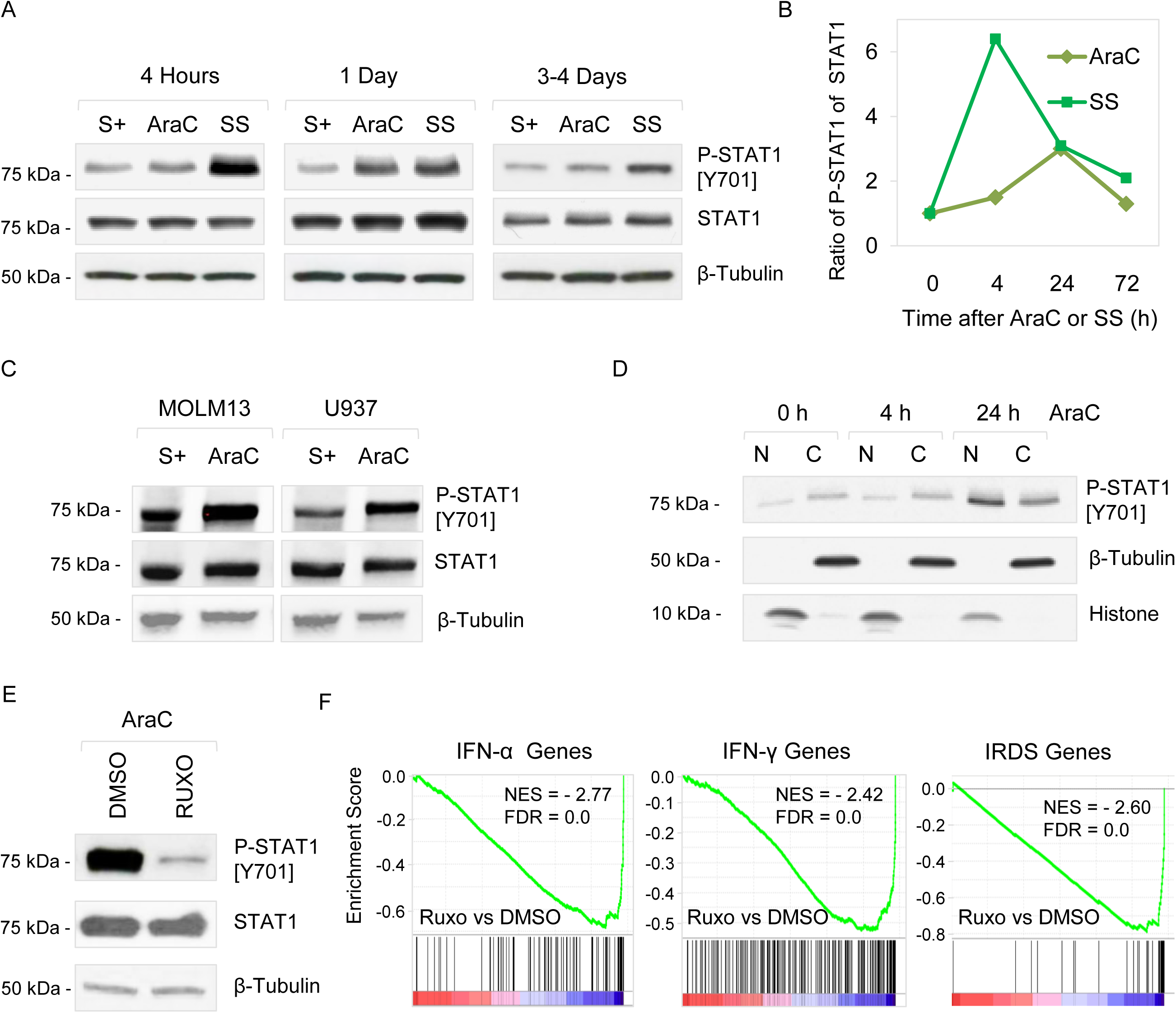
Phosphorylation of STAT1 initiates the transcription of IRDS and IFN-stimulated genes in G0 leukemic cells. **A.** Western blot analysis of phospho-STAT1 (p-STAT1-[Y701]) and STAT1 in lysates of THP1 cells treated with serum-starvation or AraC for indicated time points. **B.** The ratio of p-STAT1 to STAT1 **C.** Western blot analysis of p-STAT1 and STAT1 in different leukemic cell lines treated with AraC or vehicle for one day. **D.** Western blot analysis of p-STAT1, tubulin and Histone H3 in nuclear (N) and cytoplasmic (C) fractions of THP1 cells treated with 5 μM AraC for indicated time points. **E**. Western blot analysis of p-STAT1 and STAT1 in THP1 cells pretreated with 1 μM ruxolitinib (ruxo) or vehicle prior to AraC chemotherapy. P-STAT1-[Y701] is normalized for total STAT1 levels. β-tubulin and Actin serve as loading control. **F**. GSEA analysis of IFN-stimulated and IRDS genes, using the transcriptome of THP1 cells pretreated with 1 μM ruxolitinib or vehicle prior to AraC exposure.

### Translational down-regulation of IFN-stimulated genes in AraC-resistant leukemic cells

In order to test if IFN-stimulated genes are translationally regulated in resistant G0 leukemic cells, we examined their expression at the transcriptome and translatome levels. Overall expression of IFN-α and IFN-γ stimulated genes in G0 leukemic cells induced by serum-starvation is similar between transcriptome and translatome datasets, indicating that the increase in these conditions is due to transient transcriptional regulation (Fig. 3A). In contrast, in G0 cells induced by AraC treatment, expression of IFN-stimulated genes that is increased at 24 h of AraC treatment along with STAT1 phosphorylation (Fig. 1B, 2B), is significantly attenuated in the translatome of 3-4 days of AraC treatment, compared to RNA levels (Fig. 3A). This result suggests that the early, transient increase of IFN-stimulated genes is translationally inhibited at 3-4 days of AraC treatment. RPL13A is a component of the ribosomal subunit and plays an important role in translational inhibition of IFN stimulated genes. RPL13A forms the IFN-γ-activated inhibitor of translation (GAIT) complex [50–53] that inhibits translation of mRNAs that are transcriptionally induced by IFN-activated STAT1 pathway [54]. Based on the translational repression of IFN-stimulated genes in AraC-resistant cells, we hypothesized that RPL13A could be differentially expressed. Consistently, RPL13A is dramatically up-regulated in translatome and protein levels from 24 h through 3 days of AraC treatment. This increase was specific as most other ribosomal protein genes were down-regulated or not differentially expressed (Fig. 3B-C). Correspondingly, *VEGFA* mRNA, another target gene of the RPL13A-GAIT translational repression complex [54], is translationally attenuated in AraC-resistant cells (Fig. 3D). These data suggest that increased RPLA13A correlates with and can contribute to translation repression of IFN-stimulated mRNAs *via* the GAIT complex.

**Figure 3.**
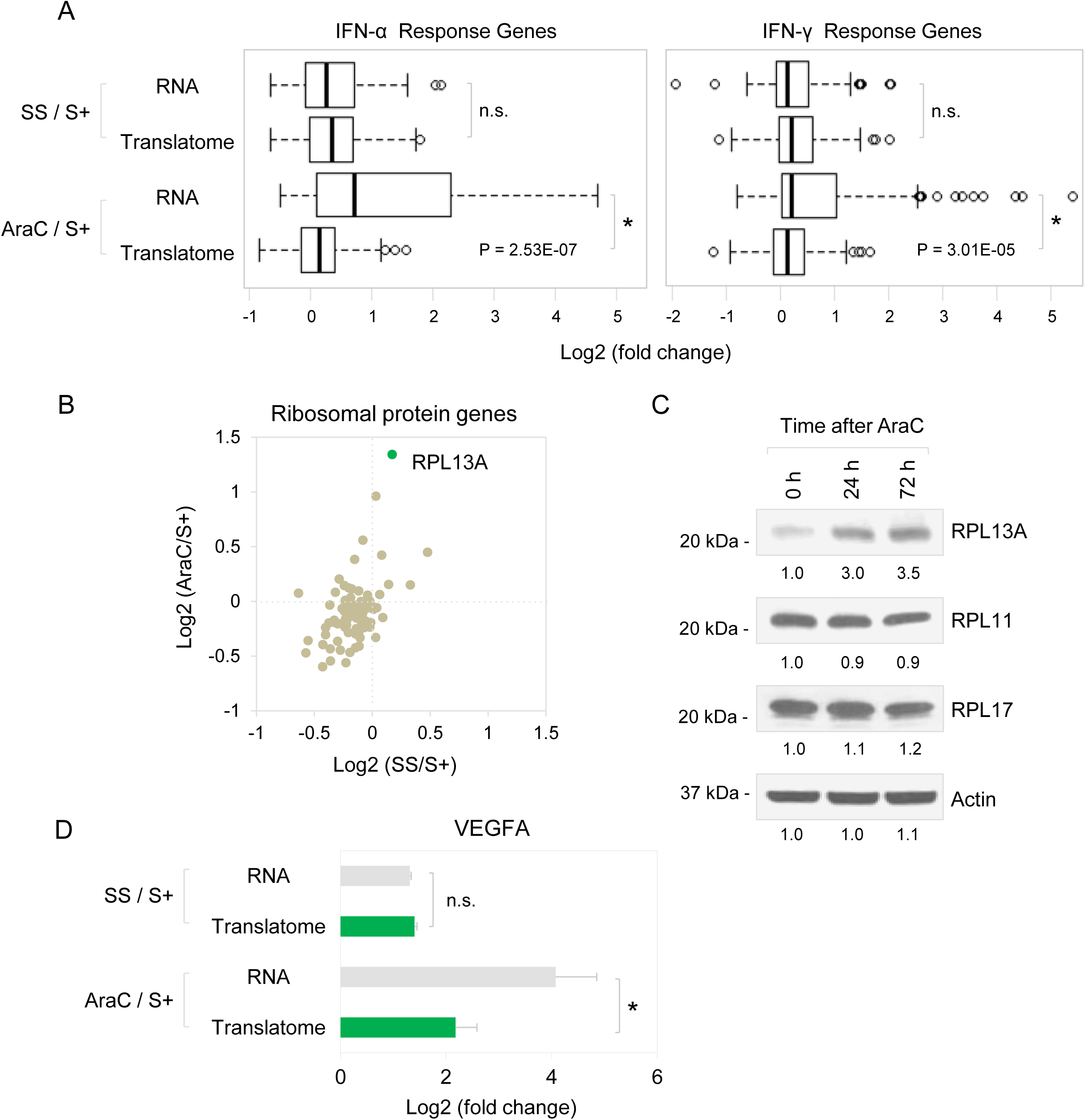
Translational regulation of gene expression in G0 leukemic cells. **A.** Expression of IFN-α and IFN-γ stimulated genes in the transcriptome and translatome levels in response to AraC or serum-starvation. **B**. Expression of ribosomal protein genes in the translatome level upon AraC or serum-starvation treatment. RPL13A is indicated with a green circle. **C**. Western blot analysis of RPL13A, RPL11, RPL17 and Actin (loading control) in THP1 cells treated with AraC for indicated time points. **D**. *VEGFA* expression at the transcriptome and translatome levels in G0 leukemic cells. \**P* ≤ 0.05. N.S. indicates not significant

### Activation of JNK and p38 MAPK is required for STAT1 phosphorylation and expression of IFN-stimulated genes

We demonstrated that STAT1 transcriptionally induces the expression of IFN-stimulated genes (Fig. 4A). To study how phosphorylation of STAT1 is regulated in resistant G0 leukemic cells, we examined MAPK signaling pathways which are activated by various cellular stresses [55–63] (Fig. 4A). As shown in our previous study [4], p38 MAPK was rapidly phosphorylated and activated within two hours of AraC treatment (Fig. 4B). Importantly, JNK MAPK was also quickly phosphorylated within few hours of AraC treatment, prior to STAT1 phosphorylation (Fig. 4B). Based on the sequential phosphorylation of these MAPKs first and then STAT1, we hypothesized that activation of JNK or p38 MAPK is required for STAT1 phosphorylation (Fig. 4A). To test this, we pharmacologically inhibited MAPK signaling pathways (Fig. 4A) and examined phosphorylation of STAT1 by Western blot analysis. As shown previously, p38 MAPK inhibitors such as LY2228820 [64–66] and BIRB796 [67], effectively blocked the p38 MAPK downstream effector, MK2, while JNK inhibitor, JNK-IN-8 [68], also effectively reduced phosphorylation of the downstream JNK target, c-JUN, in THP1 G0 cells [4]. Importantly, when either JNK or p38 MAPK was inhibited, STAT1 phosphorylation was significantly inhibited in AraC-treated THP1 cells (Fig. 4C). IFIT1, IFIT2, and ISG15 genes are key IFN-γ stimulated genes and greatly up-regulated in AraC-resistant leukemic cells (Fig. 4D). Expression of these genes is also significantly reduced by either JNK or p38 MAPK inhibition in AraC-treated THP1 cells (Fig. 4E), consistent with inhibited phosphorylation of STAT1 (Fig. 4C). Together, these results suggest that activation of JNK and p38 MAPK is required for STAT1-mediated expression of IFN-stimulated genes in AraC-resistant leukemic cells.

**Figure 4.**
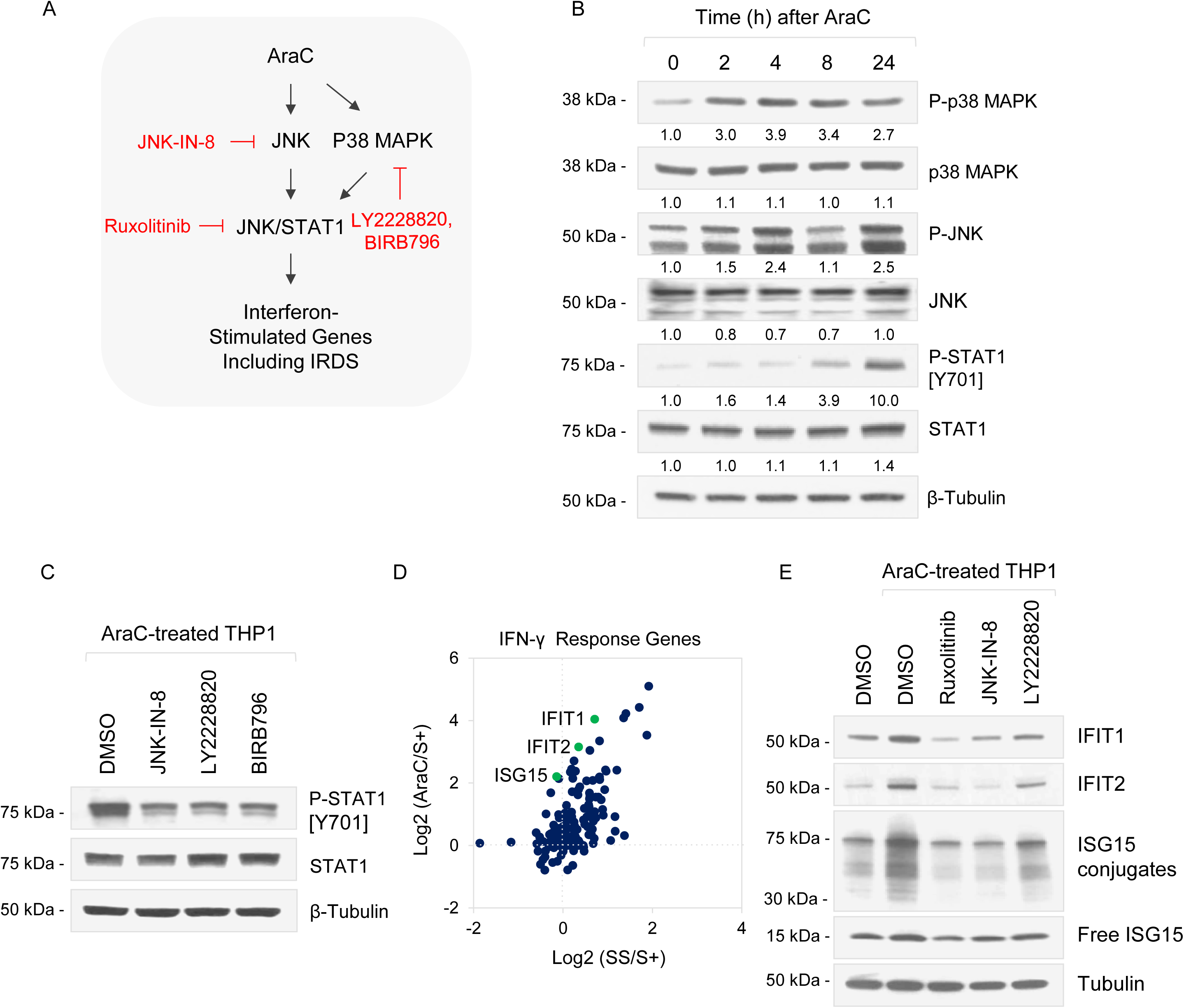
Activation of JNK and p38 MAPK are required for STAT1 phosphorylation and expression of IFN-stimulated genes. **A.** In response to AraC chemotherapy, leukemic cells activate signaling pathways such as STAT1, p38 and JNK MAPK to express IFN-stimulated genes. JAK-STAT1 signaling is inhibited by ruxolitinib, JNK MAPK by JNK-IN-8, and p38 MAPK by either LY2228820 or BIRB796. **B**. Western blot analysis of P-STAT1-[Y701], P-JNK, P-p38 MAPK, STAT1, JNK and p38 MAPK in THP1 cells treated with 5 µM AraC for indicated time points. **C**. Western blot analysis of P-STAT1-[Y701], STAT1 and Tubulin in THP1 cells pretreated with 1 µM JNK-IN-8, 5 µM LY2228820, 5 µM BIRB796 or DMSO vehicle prior to 5 µM AraC treatment. **D**. Scatter plot shows expression of IFN-γ-stimulated genes at the translatome level upon AraC or serum-starvation treatment. IFIT1, IFIT2 and ISG15 are indicated with green circles. **E**. Western blot analysis of IFIT1, IFIT2 and ISG15 in THP1 cells pretreated with 1 µM ruxolitinib, 1 µM JNK-IN-8, 5 µM LY2228820 or DMSO vehicle prior to 5 µM AraC treatment. P-STAT1-[Y701] is normalized for total STAT1 levels. β-tubulin and Actin serve as loading control.

### IFN-stimulated genes do not promote survival of AraC-resistant leukemic cells and are not associated with overall survival rates of patients with hematological malignancies

We demonstrated that JNK or p38 MAPK phosphorylates STAT1, resulting in the expression of IFN-stimulated genes (Fig. 4A). These include IRDS genes that are known to be associated with therapy resistance in solid tumors [30–42]. To investigate whether IFN-stimulated genes promote the survival of AraC-resistant cells, their expression was suppressed by JAK/STAT, JNK and p38 MAPK inhibitors, prior to AraC treatment, followed by measurement of cell viability and caspase activity to measure cell death. In our previous study, we found that p38 MAPK inhibition prior to AraC severely reduced the survival of AraC-resistant cells by destabilizing TNFα and DUSP1 mRNAs [4]. In this study, JNK inhibition with JNK-IN-8 protected leukemic cells from AraC-mediated apoptosis in THP1 and MOLM13 cell lines, and did not affect the viability of untreated cells as a control (Fig. 5A). This result suggests that in contrast to p38 MAPK which induces chemoresistance, JNK sensitizes leukemic cells to AraC treatment. On the other hand, STAT1 inhibition with ruxolitinib did not affect the viability of both AraC-treated and untreated leukemic cells (Fig. 5A). Together, these results suggest that even though activation of JNK, p38 MAPK, and STAT1 are required for the expression of IFN-stimulated genes, the expression of IFN-stimulated genes is not associated with therapy survival but likely with other cell biological roles in resistant AML cells. To test if the expression of IFN-stimulated genes affects the overall survival rates of patients with hematological malignancies, Kaplan-Meier analysis was conducted. We found that survival rates were not correlated with the expression level of IFN-stimulated genes in datasets of patients with AML, CML and lymphoma (Fig.5B), consistent with no effect of STAT1 inhibition on survival of AraC-treated cells (Fig. 5A). These data suggest that IFN-stimulated genes can promote therapy-resistance only in specific cancers rather than in hematological malignancies.

**Figure 5.**
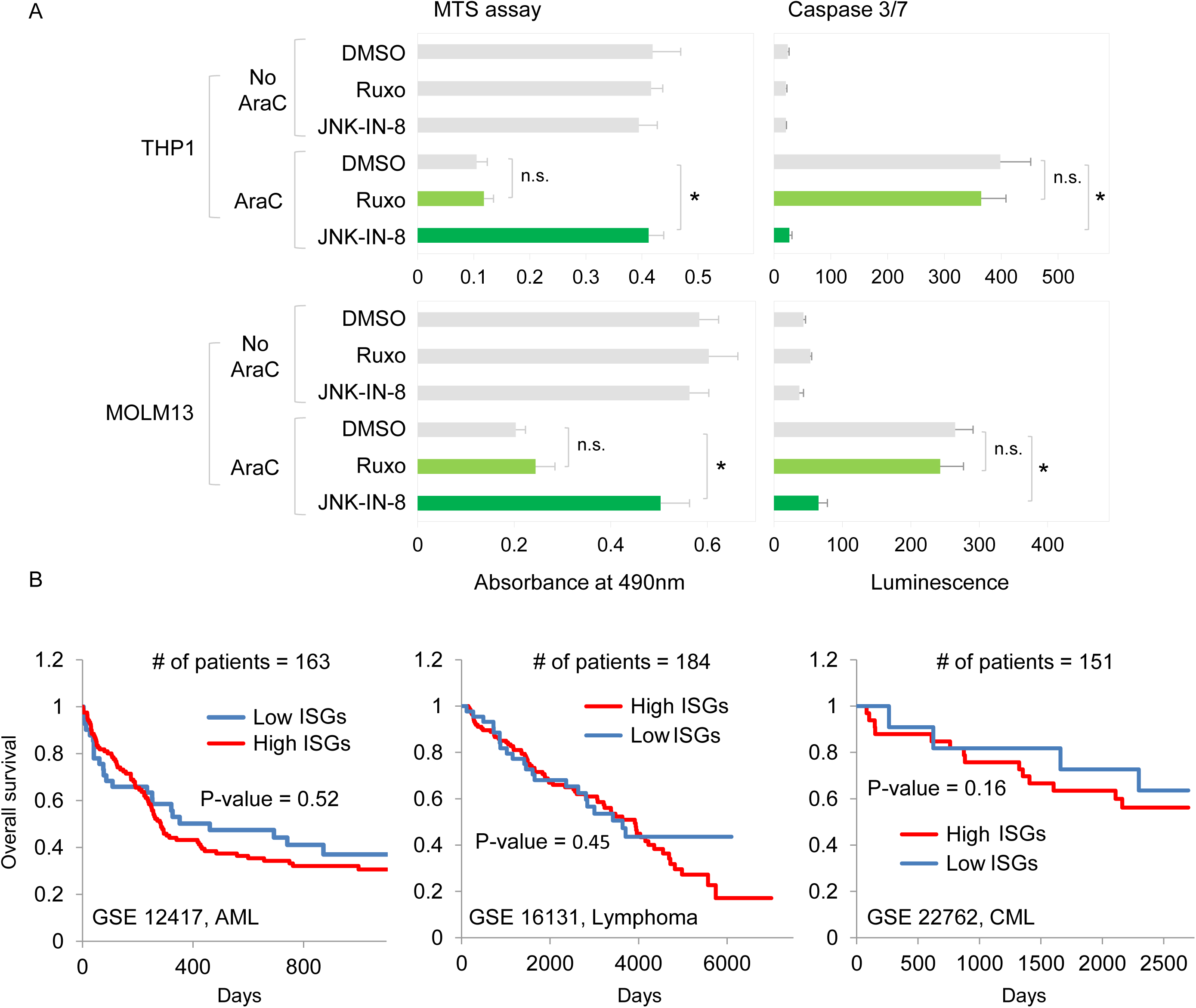
Effect of IFN-stimulated genes on chemoresistance and overall survival rates. **A.** Effect of pharmacological inhibition of IFN-stimulated genes on viability of leukemic cells. THP1 and MOLM13 cells were treated with 1 µM JNK-in-8, 1 µM ruxolitinib, and DMSO vehicle in the absence or presence of AraC treatment. After 3 days of incubation, cell viability and caspase 3/7 activity were measured. **B.** Correlation between expression of IFN-stimulated genes and overall survival rates of patients with hematological malignancies. The correlation between gene expression and overall survival rates was analyzed using PROGgeneV2 [103]. \**P* ≤ 0.05. Data are represented as average ± SEM. N.S. indicates not significant

### Resistant G0 leukemic cells show increased cell adhesion and induction of cell migration

Cell adherence and induction of cell migration play important roles in cancer invasion and metastasis [69–77]. To measure these functions in G0 leukemic cells, we used fibronectin-coated wells [78] and transwell assays [79–82] (Fig. 6A-B). Interestingly, we found that G0 leukemic cells were more adherent on fibronectin-coated plates than proliferating leukemic cells (Fig. 6A). In addition, the conditioned media from G0 leukemic cells enhanced migration of co-cultured GFP-THP1 and MCF7 cells (Fig. 6B). These data are consistent with our translatome and proteome datasets of G0 leukemic cells and other G0 cells [4] that reveal significant increase in expression of cell adhesion and cell migration inducing genes.

**Figure 6.**
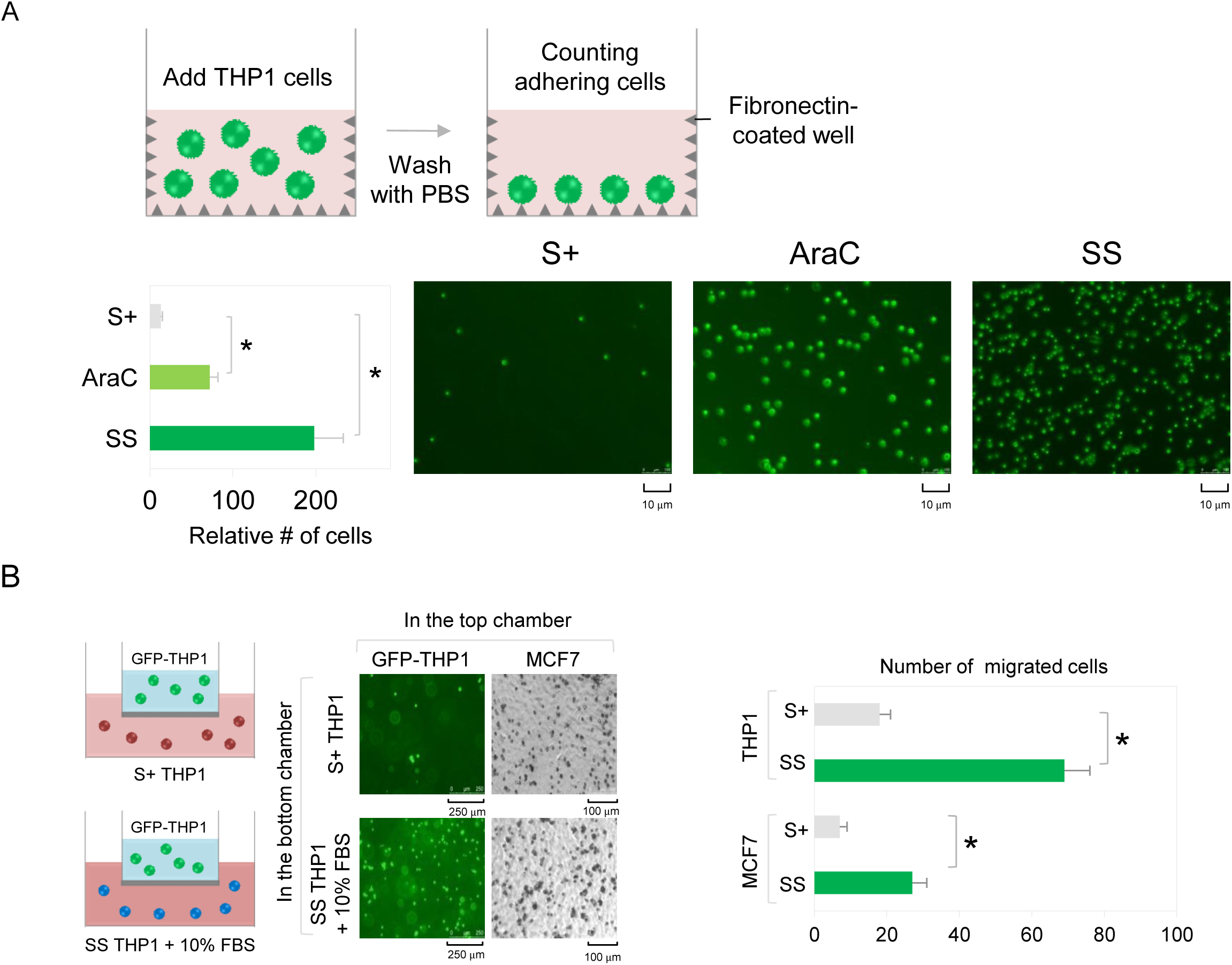
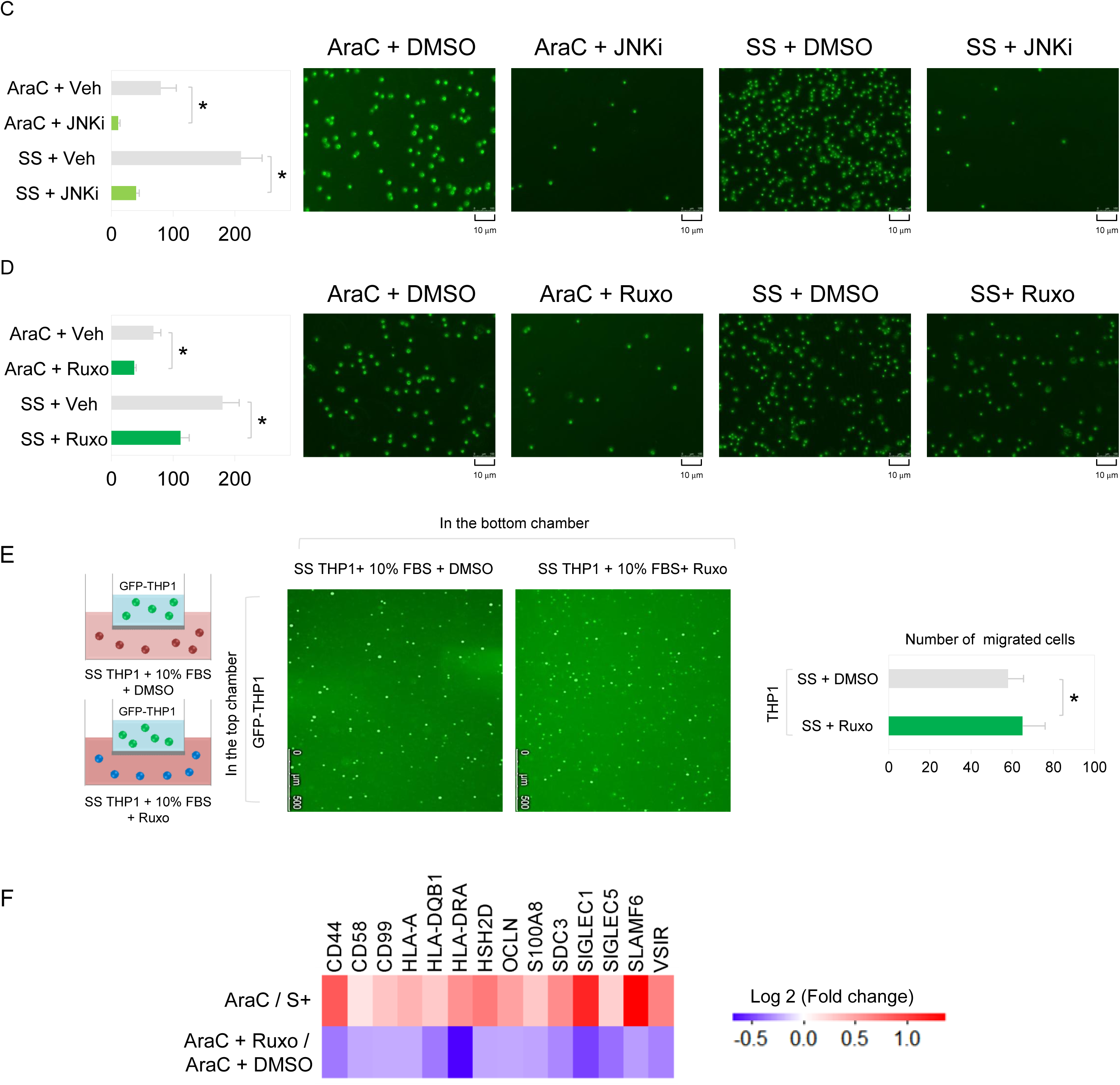
Inhibition of IFN-stimulated genes blocks adhesion of resistant G0 leukemic cells, but does not diminish cell migration of GFP-tagged THP1 monocytes or MCF7 cells. **A.** Cell adhesion assay of leukemic cells. Proliferating and G0 leukemic cells were incubated with fibronectin (FN)-coated plates for 2 hours and then washed with PBS. The number of cells bound to the coated plates was calculated and representative microscopic images are shown. **B**. Transwell migration assay. GFP-tagged THP1 cells and MCF7 cells that were plated in the top chamber were co-cultured with proliferating or G0 leukemic cells in the bottom chamber. The number of GFP-tagged THP1 or MCF7 cells that migrate through the transwell membrane to the bottom chamber is graphed and representative microscopic images are shown. **C**. Cell adhesion assay of resistant G0 leukemic cells treated with 1 µM JNK-in-8 or DMSO vehicle. **D**. Cell adhesion assay of resistant G0 leukemic cells treated with 1 µM ruxolitinib or vehicle. **E**. Transwell migration assay of GFP-tagged THP1 cells co-cultured with G0 leukemic cells treated with 1 µM ruxolitinib or vehicle. **F**. Expression of a subset of cell adhesion genes and immunoglobulin superfamily in G0 versus S+ cells and G0 cells treated 1 µM ruxolitinib versus G0 cells treated with DMSO vehicle (transcriptome). \**P* ≤ 0.05. Data are represented as average ± SEM. N.S. indicates not significant.

### The JNK-STAT1 signaling pathway promotes cell adhesion of resistant G0 leukemic cells

To measure the effect of the JNK-STAT1 pathway on adhesion and induction of migration of resistant G0 leukemic cells, the JNK-STAT1 pathway was suppressed by JNK or STAT1 inhibition. Importantly, JNK inhibition greatly reduced the number of G0 leukemic cells that were attached on fibronectin-coated plates (Fig. 6C). When STAT1 and IFN-stimulated genes were directly inhibited by ruxolitinib, the number of adherent cells also decreased (Fig. 6D), consistent with the decreased cell adhesion observed with upstream inhibition of JNK (Fig. 6C). However, STAT1 inhibition did not diminish migration of co-cultured GFP-tagged THP1 and MCF7 cells (Fig. 6E). Together, these results suggest that the JNK-STAT1 pathway promote adherence of G0 leukemic cells. These data correspond to increased expression of cell adhesion genes in G0 leukemic cells (Fig. 6F). The immunoglobulin superfamily is a group of cell surface proteins that are involved in the adhesion or recognition of cells [83–89]. Interestingly, we found that a subset of the immunoglobulin superfamily and cell adhesion genes that are highly expressed in G0 leukemic cells, are suppressed upon STAT1 inhibition (Fig. 6F), indicating that these genes are targets of IFN-STAT1 pathway, and can contribute to cell adhesion in resistant G0 leukemic cells.

## Discussion

In response to viral infection and other signaling pathway triggers, mammalian cells express diverse IFN-stimulated genes to exert antiviral functions [5, 6, 8, 9], and to control innate and adaptive immune response that impact anti-tumor immunity and the tumor microenvironment [5, 10, 12–20]. In this study, we reveal that AraC chemotherapy and serum-starvation treatment induces IFN-stimulated genes at the transcriptome, translatome, and proteome levels in acute monocytic leukemic cells (Fig.1B). Furthermore, gene signatures of *in vivo* resistant models of leukemia such as LRC [43] and LIC [44] include IFN-stimulated and immune response genes (Fig. 1D), indicating the clinical relevance of our observation. We further show that IFN-stimulated genes are transcriptionally induced by STAT1 phosphorylation in G0 leukemic cells (Fig. 2).

Our data revealed that AraC treatment or serum-starvation induced phosphorylation of STAT1 (Fig. 2A). Phosphorylation on tyrosine 701 and serine 727 of STAT1 is involved in its activation [90]. IFN binding to its receptor leads to tyrosine 701 phosphorylation of STAT1 by JAK, which is required for its localization to nucleus and enables its roles in transcription. In addition, phosphorylation on serine 727 is required. These phosphorylation events lead to full transcriptional activation of STAT1 in IFN signaling [91]. Our data reveal that AraC treatment or serum-starvation activates both p38 MAPK and JNK (Fig. 4B) [4]. Apart from the known effect of p38 MAPK on phosphorylation of serine 727 [92, 93], we find that both p38 MAPK and JNK control phosphorylation of STAT1 on tyrosine 701 (Fig. 4C), permitting full activation of STAT1-mediated transcription of IFN-stimulated genes in G0 leukemic cells.

Although previous studies have shown how phosphorylation of STAT protein is regulated by MAPK signaling [94–96] in response to LPS, UV and TNFα [97], regulation of STAT and IFN is less understood in chemotherapy-induced G0 leukemic cells. Activation of p38 MAPK was shown to enhance STAT1 phosphorylation upon UV and LPS treatment [92, 93]. In non-leukemic cells, activation of JNK with UV irradiation or its upstream kinase MKK7 has been shown to significantly block STAT3 tyrosine phosphorylation and transcription of IFN-stimulated genes [98]; however, in G0 leukemic cells, AraC treatment activated JNK but did not induce any changes in STAT3 phosphorylation (data not shown). In chemoresistant G0 leukemic cells, our data reveal that both JNK and p38 MAPK play an essential role in STAT1 phosphorylation and transcriptional expression of IFN-stimulated genes. Pharmacological inhibition of JNK or p38 MAPK drastically inhibited STAT1 phosphorylation and expression of IFN-stimulated genes (Fig. 4C-E). These data suggest that the effect of MAPKs on activation of STAT proteins is specific for cell type and environmental stimuli.

Interestingly, we observed that IFN-stimulated genes that are transcriptionally induced by STAT1 phosphorylation upon early times of AraC treatment, are subsequently, at later times, translationally down-regulated. RPL13A can translationally decrease expression of a subset of IFN-stimulated genes [53, 54]; consistently, we find that RPL13A is significantly increased in AraC-resistant cells (Fig. 3B-C). Correspondingly, RPL13A targets including *VEGFA* (Fig. 3D) and other IFN-stimulated genes are translationally reduced.

Our findings reveal that resistant G0 leukemic cells are more adherent and induce migration of neighboring immune cells (Fig. 6A-B). We demonstrate that the JNK-STAT1 pathway is responsible for adherence of resistant G0 leukemic cells but does not underlie the increased induction of cell migration observed in resistant G0 leukemic cells (Fig. 6C-E). AML cells express cell adhesion molecules on their surface to interact with the bone marrow microenvironment [99, 100] and IFNs alter their microenvironment to enable tumor persistence [19, 20]. This interaction is advantageous for tumor persistence as it protects leukemic cells from chemotherapy and promotes disease relapse [99, 100]. Therefore, although inhibitors of STAT1 or JNK do not increase chemosensitivity *in vitro* (Fig. 5A), inhibiting this JNK-STAT1 pathway could support therapeutic targeting of drug-resistant leukemic cells *in vivo* by reducing their cell adhesion ability; and thereby, curb their interaction with the bone marrow niche, which is critical for tumor survival. The JAK-STAT1 and MAPK pathways [94–96] interconnect with other cancer signaling pathways, which can impact tumor persistence in multiple ways. The JNK and p38 MAPK signaling intersect with the KRAS/MAPK pathways that impact cell cycle state and cancer metabolism [101, 102]. Given this interplay, the JNK-STAT1 pathway may impact AML chemosurvival via the KRAS/MAPK regulation of the quiescent cell cycle state. Additionally, the JNK-STAT1 pathway crosstalk with KRAS/MAPK, may influence AML persistence through regulation of cancer metabolism. Therefore, given these important signaling interactions, future studies can investigate the impact of JNK-STAT1 pathway on the quiescent cell state that is advantageous for AML chemosurvival, and on AML survival via cancer metabolism modulation, via crosstalk with important cancer signaling controllers such as KRAS/MAPK. In conclusion, our study reveals that a JNK-STAT1 pathway controls expression of IFN-stimulated genes and promotes cell adhesion in G0 chemoresistant leukemic cells.

## Acknowledgements

The study is funded by NIH grants: GM100202, CA185086, & R35GM134944 and the Brown RNA Center funds to SV. We thank past and current Vasudevan lab members for feedback and technical assistance. PASB is supported by a NIH IMSD T32 fellowship. SL was supported by MGH ECOR Fund for Medical Discovery fellowship & by a postdoctoral fellowship from the Basic Science Research Program through the National Research Foundation of Korea (2015017218).

## Contributions

SL conducted the study and helped write the manuscript. AB, PASB, EP, ZK, OC, conducted experiments for the revision. SV supervised the study and wrote the manuscript.

## Data Availability

Raw datasets are available on the public repository, as supplementary tables or on GEO as described in Lee et al., Genome Biol 2020, and all materials will be made available publicly on publication and on request.

---

**Table S1. RNA and translatome levels of IFN**-α **and IFN**-γ regulated genes in 24 h AraC treated G0 THP1 cells compared to untreated proliferating THP1 cells.

